# Multiple sequence-alignment-based RNA language model and its application to structural inference

**DOI:** 10.1101/2023.03.15.532863

**Authors:** Yikun Zhang, Mei Lang, Jiuhong Jiang, Zhiqiang Gao, Fan Xu, Thomas Litfin, Ke Chen, Jaswinder Singh, Xiansong Huang, Guoli Song, Yonghong Tian, Jian Zhan, Jie Chen, Yaoqi Zhou

## Abstract

Compared to proteins, DNA and RNA are more difficult languages to interpret because 4-letter-coded DNA/RNA sequences have less information content than 20-letter-coded protein sequences. While BERT (Bidirectional Encoder Representations from Transformers)-like language models have been developed for RNA, they are ineffective at capturing the evolutionary information from homologous sequences because unlike proteins, RNA sequences are less conserved. Here, we have developed an unsupervised Multiple sequence-alignment-based RNA language model (RNA-MSM) by utilizing homologous sequences from an automatic pipeline, RNAcmap. The resulting unsupervised, two-dimensional attention maps and one-dimensional embeddings from RNA-MSM can be directly mapped with high accuracy to 2D base pairing probabilities and 1D solvent accessibilities, respectively. Further fine-tuning led to significantly improved performance on these two downstream tasks over existing state-of-the-art techniques. We anticipate that the pre-trained RNA-MSM model can be fine-tuned on many other tasks related to RNA structure and function.

## Introduction

Three essential biomacromolecules in living organisms are DNA, RNA, and proteins, all of which are linear polymers, made of a fixed number of alphabet letters (typically 4 for DNA and RNA and 20 for proteins). The sequences of different letter combinations encode their biological functions, very much like meaningful sentences in human languages. As a result, language models such as BERT (Bidirectional Encoder Representations from Transformers)^1^ and GPT (Generative Pre-trained Transformer)^2,3^, originally developed for natural language processing, found their ways in dissecting sequence-structure-function relation of biological sequences^4,5^.

Most previous efforts have been focused on proteins. Examples are UniRep^6^, UDSMProt^7^, ESM-1b^8^, TAPE^9^, ProteinBERT^10^, and ProtTrans^11^. UniRep^6^ applied a recurrent neural network to learn unified representation of proteins and examined its ability to predict protein stability and mutation effects. UDSMProt^7^ developed a universal deep sequence model based on Long Short-Term Memory (LSTM) cells and demonstrated it by using several classification tasks, including enzyme class prediction, gene ontology prediction, and remote homology detection. ESM-1b^8^ is a deep transformer model trained on 250 million protein sequences and tested on secondary structure, tertiary contact, and mutation-effect prediction. TAPE^12^ systematically evaluates different self-supervised learning methods on protein sequences by utilizing five downstream tasks. ProteinBert^10^ combined a bidirectional language modeling task with a Gene Ontology annotation prediction task to pretrain a protein sequence language model. Its performance was evaluated on nine downstream tasks associated with protein structure, post translational modifications and biophysical attributes. ProtTrans^11^ compared six natural language models (including Transformer-XL, XLNet, BERT, Albert, Electra, T5) pretrained on UniRef^13^ and BFD^14^ protein sequence databases and suggested that the embedding of these protein language models have learned some “grammar” from protein sequences.

More recent attentions have been on DNAs and RNAs. These include Bert-like models for DNA, including DNABert^15^ with downstream prediction task of promoters, splice sites and transcription factor binding sites, iEnhancer-BERT^16^ with a focus on enhancer identification, and BERT6mA^17^ for predicting N6-methyladenine sites. For RNAs, preMLI^18^ trained a rna2vec model^19^ to obtain the RNA word vector representation for predicting microRNA–lncRNA interactions. RNA-FM^20^ constructed a BERT-based RNA foundation model on large unannotated RNA sequences with applications on prediction of structural and functional properties.

However, unlike human languages, whose evolution information is mostly lost in history, evolutionary history of biological sequences is well preserved through the appearance of proteins (or DNAs/RNAs) with the same function but different sequences in different species. Such evolutionary history of sequences, revealed from multiple sequence alignment (MSA), allows an in-depth analysis of sequence conservation and mutational coupling due to structural and functional requirements ^[21–23]^. Indeed, employing homologous sequences was one of the key factors for the recent success of AlphaFold2^14^ for highly accurate prediction of protein structures. Similarly, incorporating evolutionary information in a language model (MSA transformer) allows improved performance in protein contact-map and secondary-structure prediction over a single-sequence-based language model with fewer parameters^24^.

To build a similar MSA transformer for RNA, one must obtain RNA homologs first. Searching RNA homologs, however, is more challenging than searching for protein homologs due to the isosteric nature of RNA base pairs and because sequence identity is much easier to lose for a sequence with fewer alphabet letters (4 versus 20). A sequence-based search such as BLAST-N^25^ against a nucleotide database often leads to a few homologs if any. A more sensitive approach is to search for homologous sequences compatible with a defined secondary structure because secondary structure is more conserved than sequence^26^. Currently the best secondary-structure-based approach ^27,28^ is Infernal^29^, which is based on a covariance model. Infernal was employed to build RNA families (Rfam) by manual curation with known or predicted experimental secondary structures^30^. However, Rfam updates slowly and only about 4000 families have been curated so far. Moreover, using experimental secondary structure for alignment in some RNAs prohibits the application of Rfam MSAs to “ab initio” structure prediction. This problem was solved with the development of RNAcmap^31^, which integrates BLAST-N, Infernal, and a secondary structure predictor such as RNAfold^32^ for a fully automatic homology search. RNAcmap was further improved with additional iteration^33^ and a large expansion of sequence database^34^.

This work reports an RNA MSA-transformer language model (RNA-MSM) based on homologous sequences generated from RNAcmap3. We examined the usefulness of a language model by predicting base pairs and solvent accessibility derived from three-dimensional RNA structures. Using base pairs derived from RNA structures is necessary because experimentally determined RNA structures are the only gold standard dataset for base pair annotations. A significant portion of commonly used benchmark databases of secondary structure such as RNA STRAND^35^, ArchiveII^36^, Rfam^30^, and bpRNA^37^ contained the results from comparative analysis, which may not be accurate or complete^38^. SPOT-RNA was the first end-to-end prediction of RNA secondary structure and tertiary base pairs by deep learning^38^. Although a few deep learning methods have been published since then [e.g. DMfold^39^, MXfold2^40^ and Ufold^41^], SPOT-RNA and its latest improvement with evolutionary information (SPOT-RNA2^42^) remain the only two methods trained and tested with RNA-structure-derived base pairs. For solvent accessibility, three methods RNAsnap^43^, RNAsol^44^, and RNAsnap2^45^ were developed. The first method is based on support vector machines, whereas the latter two are deep learning techniques, all with evolutionary profiles as input. Here, we showed that RNA-MSM can bring significant improvement in RNA secondary structure and solvent accessibility prediction, despite previous methods employing evolution and co-evolution information as well.

## Results

### RNA-MSM model

The MSA Transformer^24^ originally developed for MSAs of homologous protein sequences was modified for MSAs of RNA sequences. The network architecture of RNA-MSM, along with its downstream tasks is illustrated in Figure 1. Specifically, the MSA generated from the RNA query sequence by RNAcmap3 was used as an input for the network. We did not employ Rfam MSAs because the median number of sequences in Rfam families is only 45. Instead, we employed representative sequences from 3932 Rfam families, excluding those with experimentally determined structures (see Methods for more details). The two-dimensional input passes through the embedding layer and the series of axial transformer modules, where the attention operates over both rows and columns of the input. The final outputs are one-dimensional MSA sequence representation and two-dimensional residue attention map that were employed as the input for downstream tasks. The model was trained by the masked language modeling objective function (see Methods for more details).

**Figure 1.**
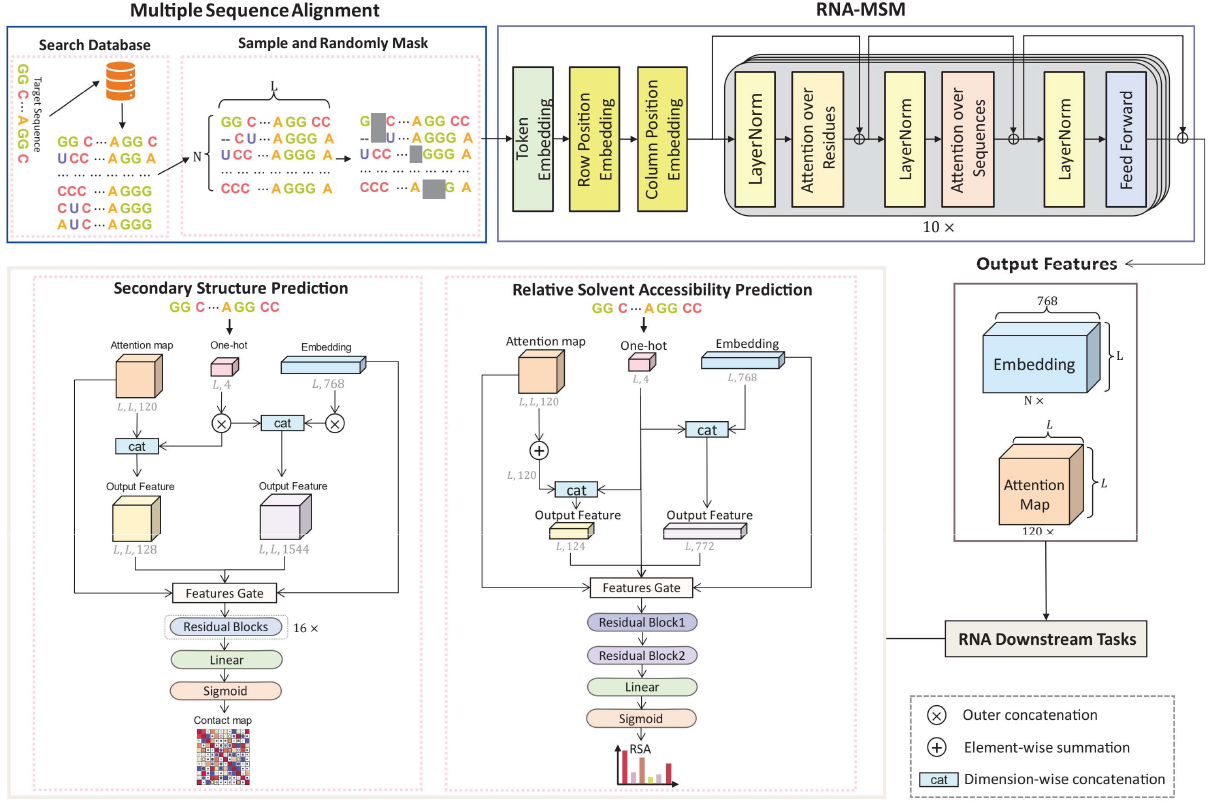
The network architecture of RNA-MSM with downstream tasks for predicting secondary structure and RNA solvent accessibility for one RNA sequence of length *L* along with *N-1* homologous sequences. The RNA-MSM model is stacked with 10 transformer blocks, each containing 12 attention heads. The output embedding identifies all sequences in a 768-dimensional space. 120 attentional maps were constructed by stitching the attention scores among the residues learned by the 12 attention heads. The feature gate used in the downstream tasks means a feature combination selection gate.

### Direct base pairing information in attention map

With the self-attention mechanism of the transformer model, pairwise interactions can be established between any positions within the sequence. In principle, multiple attention heads employed in the attention layers of transformer module should capture a variety of features from the input sequence by focusing on different portions of the input sequence simultaneously as demonstrated for proteins^46^. Here, we examined if the attention maps derived from the RNA-MSA transformer module can serve as a base-pairing probability map in the RNA sequence, using an independent TS test set.

Figure 2 shows that some attention maps can be directly employed for predicting secondary structure with reasonable accuracy (the average harmonic mean of precision and recall, F1-score, all larger than 0.5). Furthermore, top-K attention maps were selected by F1-scores and combined to predict the RNA secondary structure even more accurately. For example, Top-2 attention maps leads to an impressive performance with F1-score at 0.563, and the Matthews correlation coefficient (MCC) at 0.562, respectively (Supplementary Table S1). These unsupervised results provided the evidence that some structural information was captured by some layers in the attention maps.

**Figure 2.**
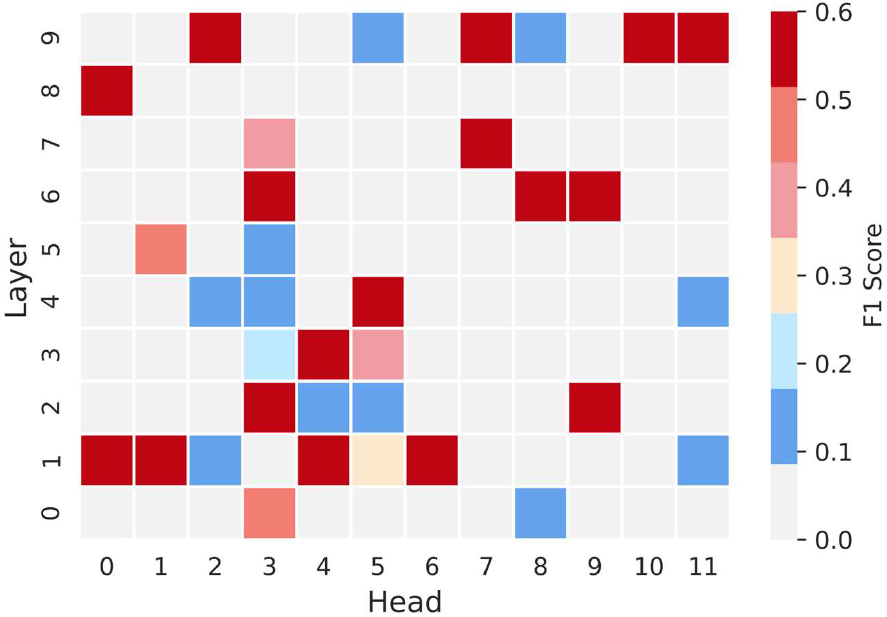
The normalized F1 scores of 120 attention maps from different layers and attention heads of RNA-MSM as input of RNA secondary structure prediction on the independent TS test set.

### RNA secondary structure prediction

#### Model Training

To further improve the secondary structure prediction beyond unsupervised learning, secondary structure prediction models were trained, validated, and tested on the secondary structure data set derived from known three-dimensional RNA structures. Briefly, the validation and test sets were structurally different (TM-Score<0.45) from each other and from the training set. In addition, the sequences in the test and validation sets were refined with sequence identity of <80% within each set. The training (TR1), validation (VL1), and test (TS) sets have 405, 40, and 70 RNAs, respectively. The secondary structure prediction model was designed based on the ResNet model as shown in Figure S1 (Details can be found in Methods).

#### Feature Comparison

To illustrate the usefulness of the features from the language models, we compared it with various previously employed features for secondary structure prediction. These features include one-hot encoding (OH), contact maps from direct coupling analysis (DCA) by gremlin^47^, sequence profiles from multiple sequence alignment (SeqProf)^29^, embedding from the RNA foundation model (RNA-FM_Emb)^20^, the logistic regression of the attention map (RNA-MSM-ATN-LR), embedding (RNA-MSM_Emb) and attention map (RNA-MSM_ATN) from our RNA-MSM model. Some feature combinations were also examined. Figure 3A compares precision-recall curves of different input feature by using the same network on the independent test set TS. One-hot encoding can be considered as the baseline model for all features compared. It is a bit surprising that the RNA-FM_Emb alone or RNA-FM_Emb combined with one hot-encoding was even worse than one hot encoding alone when employed as the features for secondary structure prediction. The AUC_PR values for the TS set are 0.340 for RNA-FM_Emb, 0.334 for OH+RNA-FM_Emb, and 0.377 for OH, respectively (Supplementary Table S2). By comparison, both sequence profile (SeqProf) and direct coupling analysis (DCA) provide boost in model performance when combined with OH with AUC_PR values at 0.377 for OH, 0.438 (OH +SeqProf), 0.501 for OH+DCA, 0.535 for OH+SeqProf+DCA, respectively (Figure 3A and Supplementary Table S2). The same is true for the features from the current model (RNA-MSM_Emb and RNA-MSM_ATN). Combining them with OH leads to AUC_PR values of 0.547 and 0.610, respectively. That is, combining one-hot encoding with the attention map from RNA-MSM yields the best model for secondary structure prediction. Interestingly, the simple logistic regression training on attention maps can achieve AUC_PR at 0.516, only worse than the models trained with attention maps (RNA-MSM_ATN) or embedding (RNA-MSA_Emb). Model performance can also be evaluated according to F1-score and Mathews Correlation Coefficients (MCC, 1 for perfect prediction and 0 for random prediction). The overall trends are the same (Supplementary Table S2). Figure 3B further shows the distributions of F1-score for each predicted RNA secondary structure by different features. The model based on OH+RNA-MSM_ATN not only has the best average F1-score, but also have the narrowest distribution, with few poor predictions. This indicates that the model based on OH + RNA-MSM_ATN has the best performance for the majority of RNAs.

**Figure 3.**
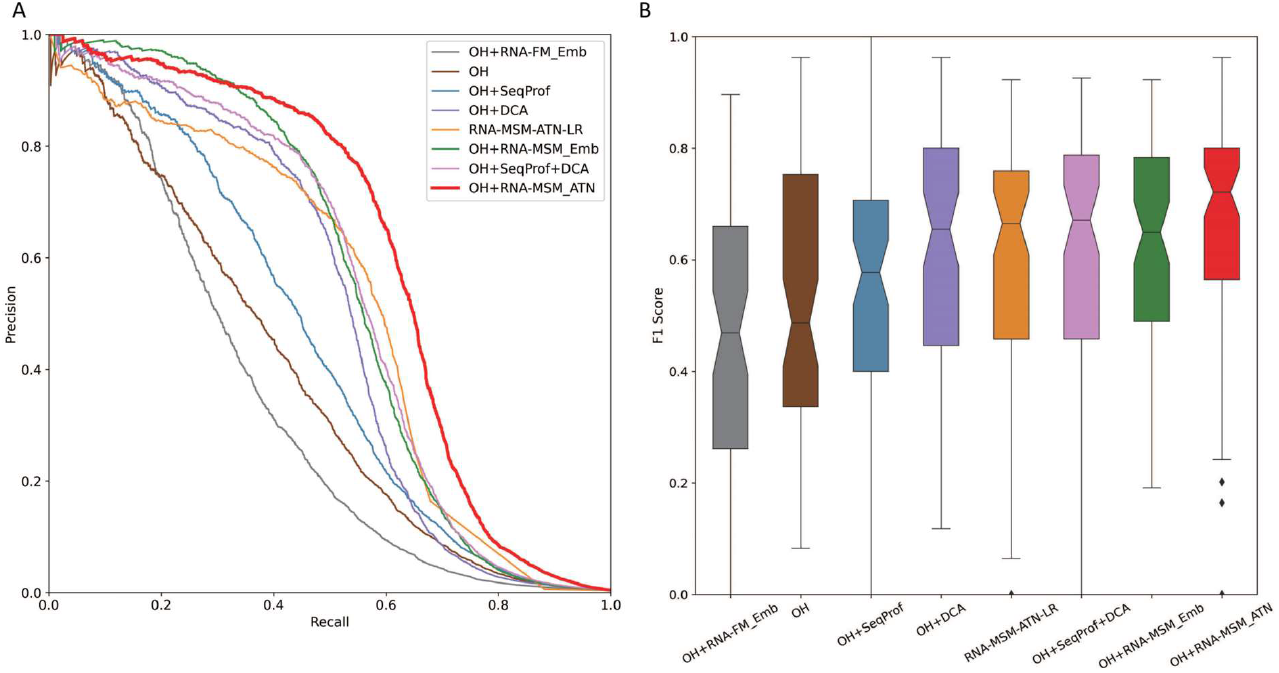
Performance comparison on the test set (TS) given various features or feature combinations according to precision-recall curve (A) and the distribution of F1-scores (B) for each RNA given by various features all trained with the same model network (Supplementary Figure S1). The features shown here are one-hot encoding (OH), sequence profiles generated by multiple sequence alignment (SeqProf), direct coupling analysis of covariation by gremlin (DCA), the embedding of RNA foundation model (RNA-FM_Emb), the logistic regression of the attention map (RNA-MSM-ATN-LR), the embedding and attention map from this work (RNA-MSM_Emb and RNA-MSM_ATN). The best model is combination of (OH+RNA-MSM_ATN) based on either area under the precision-recall curve (A) or F1 score (B).

#### Comparison to traditional methods

We compared our best secondary structure predictor (OH+RNA-MSM_ATN) with traditional folding-based single (RNAfold^48^ and linearPartitition^49^) and multi-sequence-based (CentroidAlifold^50^) techniques in Figure 4A-B and Supplementary Table 2. The alignment-based CentroidAlifold with the CONTRAfold inference engine (IE) and a gamma value of 16 were employed for its best performance as before^42^. According to the PR curves, OH+RNA-MSM_ATN has the best performance in AUC_PR (Figure 4A) and in the distribution of F1-scores (Figure 4B) for the TS set, as well as the highest F1 score and MCC value (Supplementary Table S2).

**Figure 4.**
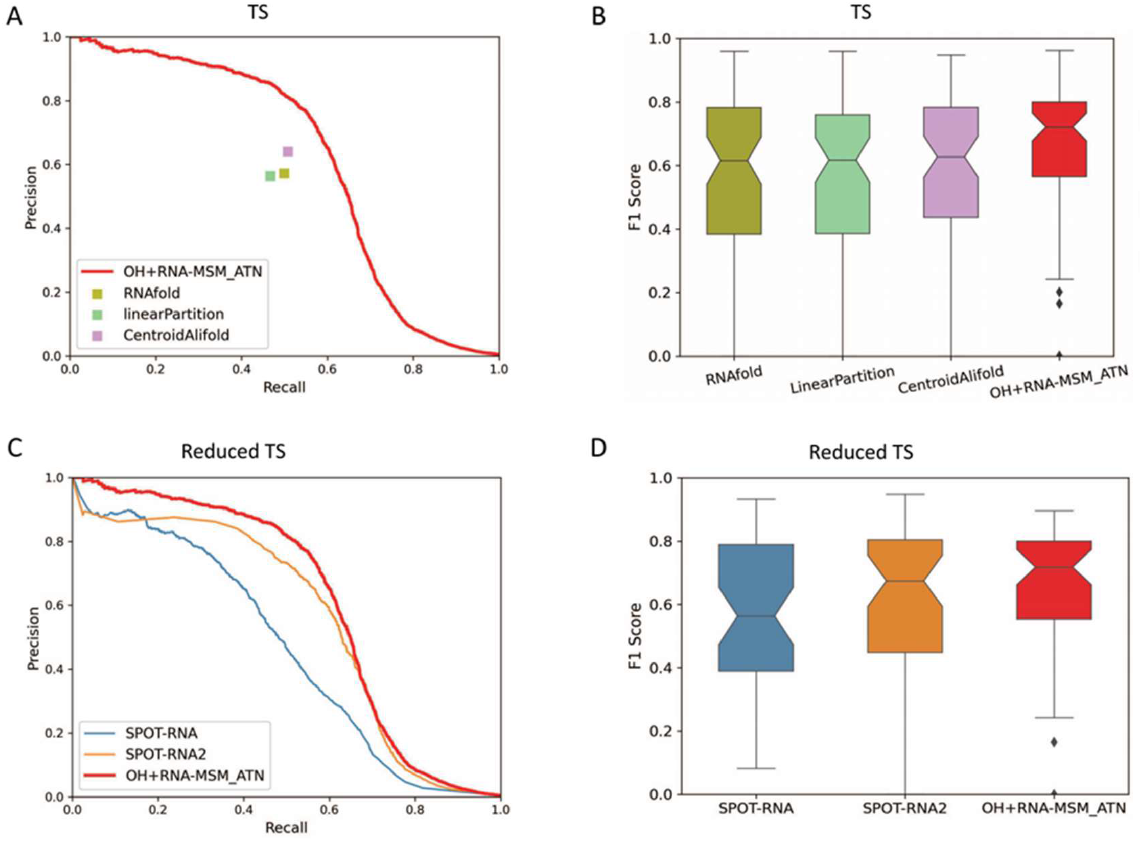
**(A)** Comparison of precision-recall curves given by OH+RNA-MSM_ATN, RNAfold, LinearPartition and CentroidAlifold on the TS test set. (B) Comparison of F1-score distributions given by OH+RNA-MSM_ATN, RNAfold, linearPartition and CentroidAlifold. **(C)** Comparison of precision-recall curves given by OH+RNA-MSM_ATN, SPOT-RNA and SPOT-RNA2 using a reduced TS set after excluding similar structures to the training set in SPOT-RNA/SPOT-RNA2. **(D)** Comparison of F1-score distributions given by OH+RNA-MSM_ATN, SPOT-RNA, and SPOT-RNA2.

#### Comparison to SPOT-RNA and SPOT-RNA2

Figure 4C compares with the only two other RNA-structure-trained secondary structure predictors (SPOT-RNA and SPOT-RNA2) using a reduced test set (48 RNAs, after removing those RNA structures similar (TM-Score>0.45) to the training set and validate set employed in SPOT-RNA and SPOT-RNA2). According to the PR curves, SPOT-RNA2 has the second-best performance with AUC_PR at 0.56 among the existing methods compared (Supplementary Table S2), compared to 0.6 by OH+RNA-MSM_ATN for the reduced set. The conclusion is also true for the distribution of F1-scores (Figure 4D). Supplementary Table S2 provides a detailed comparison for AUC_PR, F1-scores, and MCC. Here, we did not attempt to compare with other deep-learning techniques because they were not trained by 3D-structure-derived base pairs.

#### The factors controlled the performance

The wide distribution of F1-score shown in Figures 3-4 indicates that the model OH+RNA-MSM_ATN perform well on some RNAs but not others. To provide a better understanding, we examined the dependence of F1-scores on the sequence length (Supplementary Figure S2A) and on the performance of RNAfold (Supplementary Figure S2B), because RNAfold was employed as the initial secondary structure for homologous sequence search in RNAcmap3. We did not find any significant correlation between the F1-scores and sequence lengths. However, there is a strong correlation between F1-scores given by OH+RNA-MSM_ATN and that given by RNAfold with Pearson’s correlation coefficient of 0.774. Despite the influence from RNAfold prediction, most F1-scores given by OH+RNA-MSM_ATN improve over those from RNAfold (Supplementary Figure S2B), except for those that already achieved high performance (F1-score >0.7 by RNAfold).

#### Base pairs due to tertiary interactions

Supplementary Table S3 further examines the performance of different methods on canonical and non-canonical base pairs according to F1-score, precision, and recall. It shows that the method based on the MSM-language model remains the best for both canonical and noncanonical base pairs. Although predicting noncanonical base pairs are challenging, 82% improvement in F1-score over one-hot encoding is observed. When comparing to SPOT-RNA and SPOT-RNA2 for the reduced independent test set, the model OH+RNA-MSM_ATN improves over SPOT-RNA by 28% and SPOT-RNA2 by 7% in F1-score for canonical base pairs and over SPOT-RNA by 136% and SPOT-RNA2 by 4% in F1-score for noncanonical base pairs. Supplementary Table S4 further compares the performance on pseudoknots, lone base pairs and triplets. Interesting, OH+RNA-MSM-ATN is the best for lone-pairs but OH+SeqProf+DCA (F1-score=0.278) or SPOT-RNA (F1-score=0.274 for the reduced TS set) is the best for pseudoknots whereas F1-score=0.243 for OH+RNA-MSM_ATN (F1-score=0.176 for the reduced TS set).

#### Ensemble Model

An ensemble of four model (OH+SeqProf+DCA, OH+RNA-MSM_Emb, RNA-MSM_ATN, and OH+RNA-MSM_ATN) can improve the performance over OH+RNA-MSM_ATN by another 2% in F1-score. The improvement for canonical, noncanonical and pseudoknots base pairs is shown in Supplementary Table S5. The largest improvement is on pseudoknots base pairs, which now has the best performance even on the reduced TS set, when compared to SPOT-RNA and SPOT-RNA2 (Supplementary Table S4).

#### Direct solvent accessibility information in embedding

Given the fact that unsupervised attention maps contain base pairing information (Figure 2), it is of interest to examine if the RNA-MSM model can capture solvent accessibility directly from the original MSA without supervised training. To answer this question, we employed the original embeddings of RNA-MSM for this analysis. The embedding represents the MSA of a target sequence as an L × 768 matrices where L is denoted as the length of RNA chain sequence. Each nucleotide is represented by a vector of 768 channels. Here we examined the correlation between weights in channels with relative solvent accessibility (RSA) based on Spearman’s rank correlation coefficient (SCC).

Figure 5A shows the sorted mean SCC values for the TS set for different embedding channels. The result shows that the highest positive SCC score is 0.380 and the lowest SCC score is −0.427. The distribution of correlation coefficients shown in Figure 5B further shows that there are quite a few channels with weak correlations to RSA, indicating that some embedding channels did capture the structural information of RNAs without training.

**Figure 5.**
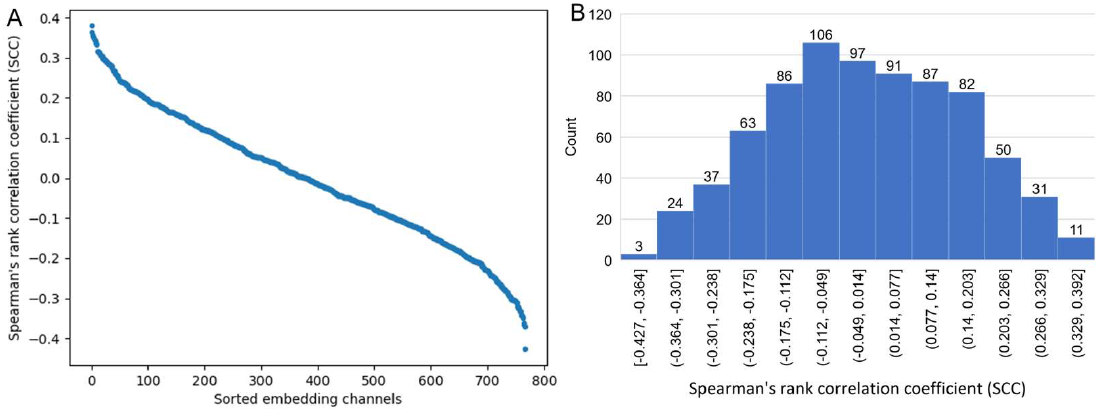
(A) The mean Spearman Correlation Coefficients (SCC, Y-axis) on the test set of 768 embedding channels (X-axis) from last layers of RNA-MSM, sorted according to the SCC values. (B) The distribution of SCC values.

#### RNA solvent accessibility prediction

The training, validation and test sets for secondary structure predictions were employed for solvent accessibility prediction. The metrics for performance of different predictors are the same as those used in RNAsnap2^45^, including PCC of individual chains between predicted and actual RSA values and mean absolute error (MAE) of individual chains between predicted and actual ASA values. Supplementary Table S6 summarizes the performance comparison among previous work and different feature combinations. One-hot encoding (OH) alone yields a reasonable performance with PCC=0.447 and MAE of 34.05 on TS. Adding the sequence profile generated from multiple-sequence alignment further improves the performance. Surprisingly, the embedding from a previous language model RNA-FM performed even worse than one-hot encoding. This is also true for the attention map from RNA-MSM. RNA-MSM_ATN alone as a feature does not do well as one-hot encoding, but its combination with one-hot-encoding is useful for going beyond one-hot-encoding. The best performing method is based on the RNA-MSM embedding (RNA-MSM_Emb). Its combination of one-hot encoding yields the lowest MAE and the highest PCC. RNA-MSM_Emb has a slightly lower PCC value and worse MAE than OH+RNA-MSM_Emb. Thus, we will choose OH+RNA-MSM_Emb as our final model for ASA prediction. The performance of the OH+RNA-MSM embedding alone is 7% improvement in PCC and 3% in MAE over the previously developed method RNAsnap2_pro, which is based on sequence profile information. The improvement is statistically significant according to the distribution of PCC and MAE values for individual RNAs (Supplementary Figure S3A-B) and p-values (Supplementary Figure S3C-F). If we removed similar structures to the sequences in the training and validating set from RNAsnap2 (the reduced set TS*), the improvement of the model OH+RNA-MSM_Emb over RNAsnap2_pro is even higher (16% improvement in PCC and 6% in MAE as shown in Supplement Table S7).

We further examined the usefulness of an ensemble model by simply averaging the outputs of individual models^51^. We test two strategies for the ensemble. The Ensemble_T model is made of the top 3 models produced by the training stage according to the PCC on VL1. The Ensemble_F model is made of one-hot-encoding (OH), RNA-MSM_Emb, and OH+RNA-MSM_Emb. As shown in Supplementary Table S7, both ensemble models show improved performance on the testing set with Ensemble_F has a slight edge. We found that Ensemble_F yielded a PCC of 0.504 and an MAE of 33.12 for the TS set. This is a 3% improvement in PCC and a 3% improvement in MAE over the best single model OH+RNA_MSM_Emb. Adding additional models only makes a minor improvement. The Ensemble_F model has a narrower distribution of PCC and MAE on the TS set than the single model (Supplementary Figure S5).

The MAE of different RNA chains diverse in a wide range when predicting their ASA, this phenomenon may be controlled by some factors. The length of RNA chains may affect the performance of the predictor. In addition, RNAfold was employed to search and generate homologous sequences by RNAcmap3. Thus, we examined the relation between ASA prediction performance and F1-score of RNAfold. We found that the PCC has a negative correlation with RNA length (Supplementary Figure S4A) but positive correlation with RNAfold F1-score (Supplementary Figure S4B).

#### Generalization beyond families trained

One interesting question is whether or not the prediction performance is robust for those families that are not in training or validation sets. Here, we examined the Rfam families in the TS set that were not in both the training set and validation set. There was a total of 8 families. Supplementary Table S8 shows the results of RNAs in these 8 families (RF01998, RF00080, RF00075, RF00028, RF01786, RF01084, RF00061, RF01051). For secondary structure (SS) prediction, the majority (6/8) has a high prediction accuracy (F1-score >0.64). The ones with a poor SS prediction have a poor RNA-fold prediction (F1-score by RNAfold <0.26A). For ASA, all have MAE values that are lower than the average performance (34.03). This result indicates that the method developed here can be generalized to the family outside the training and validation sets.

Figure 6 shows one example with the average performance of the secondary structure prediction by model OH+RNA-MSM_ATN (Figure 6C) and ASA predictor by model OH+RNA-MSM_Emb (Figure 6B) for bacterial SRP Alu domain (PDB 4WFM, Chain A). As shown in Figure 6, this RNA has a well-folded structure with a unique topology. The predicted ASA has a PCC value of 0.44 with MAE=31.52, whereas the predicted secondary structure captured most inner base pairs, including the pseudoknots. The main missing helix strand is the long-range contacts for │i-j│>87. The F1-score is 0.74.

**Figure 6.**
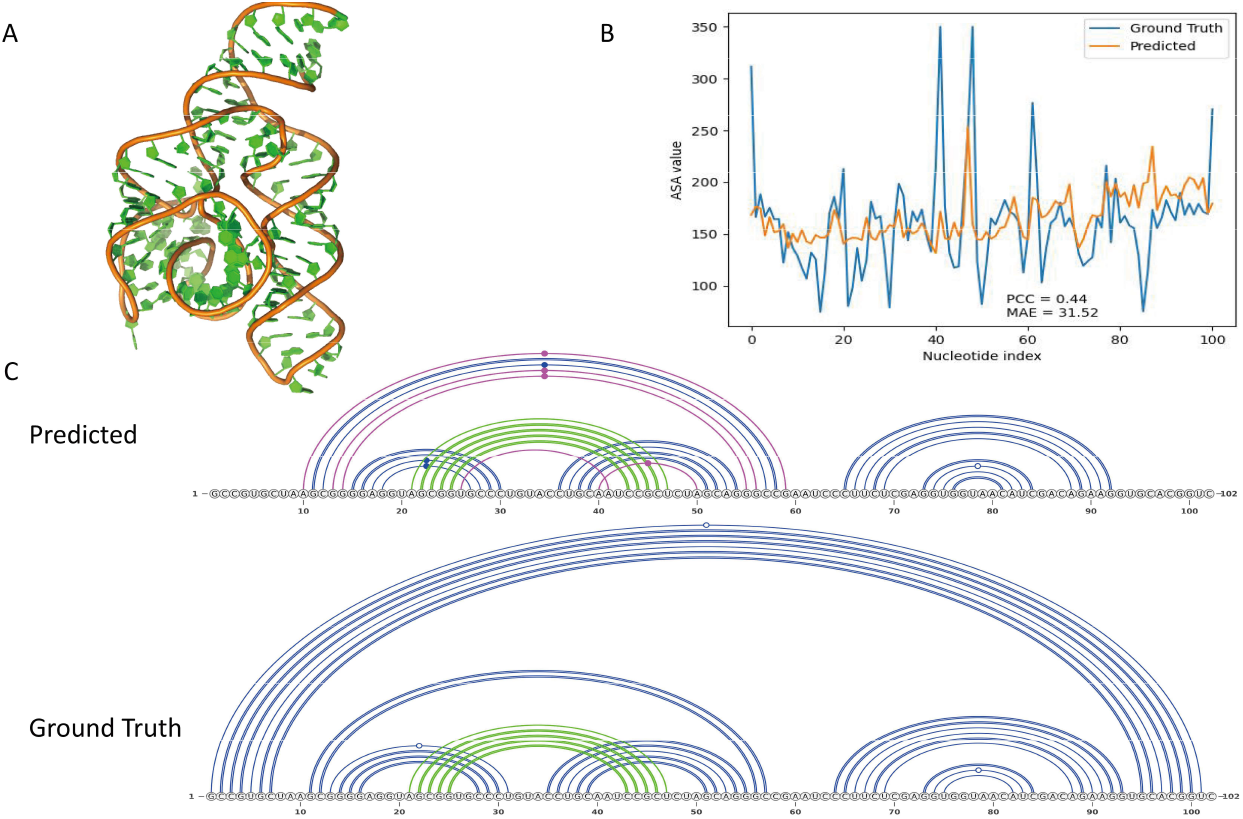
One example (chain A in PDB ID 4WFM) for ASA and secondary structure prediction with the average performance. (A) The 3D structure of 4WFM_A. (B) The result of ASA prediction. The blue line denotes the actual ASA values calculated from the native structure. The yellow line denotes the ASA values predicted by the model OH+RNA-MSM_Emb. (C). The base pairs of 4WFM_A represented by arc diagrams with canonical base pair (BP) in blue color, pseduoknot BP in green color, and wrongly predicted BP in magenta color (false positives). The predicted secondary structure by model OH+RNA-MSM_ATN with F1-Score 0.714, as compared with the native base-pairing structure for 4WFM_A RNA (Ground Truth).

## Discussion

We presented an RNA MSA-transformer language model (RNA-MSM), based on homologous RNA sequences. RNA-MSM takes the multiple aligned sequences as an input, and outputs corresponding embeddings and attention maps. We have demonstrated that these outputs contain the structure information of the query sequence of the input MSA and evaluated our model on two structure-related downstream tasks including RNA secondary structure prediction and RNA solvent accessibility prediction. Interestingly, we found that different downstream tasks prefer to different representation features. Specifically, the attention maps outperformed embeddings for RNA secondary structure prediction, while the opposite is true for the RNA ASA prediction task. It is likely that the attention maps of the two-dimensional form contain the information on the connections between nucleotides as demonstrated in Figure 2, whereas the embedding compressed the structural information into one dimension, which is more suitable for one-dimensional structural properties such as ASA.

We employed RNAcmap3 to search and generate homologous sequences. Our original plan was to use Rfam families^30^ but the median number of sequences in an Rfam family (45) is too low. Moreover, some RNA families relied on experimentally determined secondary structure for seed alignment, which would interfere with our goal to generate MSAs in the absence of any known experimental information. Thus, we employed RNAcmap3, which produced many more homologous sequences by including not only the nucleotide sequences in NCBI, but also those from RNAcentral^52^, the genomic sequences from Genome Warehouse (GWH)^53^ and the genomic sequences from MGnify^54^. Indeed, the median value for the number of MSA sequences for Rfam families expands to >2000. Significantly more homologous sequences allow a better training of the present RNA-MSM model. However, one limitation of RNAcmap3 is the time-consuming nature of obtaining the MSAs. It takes on average 9 hours for getting an MSA for one RNA of length 60. Moreover, the method needs a secondary structure predictor to provide an initial secondary structure for building the covariance model. RNAfold was used here. This leads to the overall performance somewhat dependent on the performance of RNAfold. Further studies are required to avoid the influence of the use of an initial secondary structure predictor.

Previous methods RNAsnap2_seq and RNAsnap2_pro made use of RNA secondary structure (SS) as a feature to input. We also examined the possibility of using predicted SS as an additional feature for improving RNA-MSM-based models. We did find that some improvement was made when combining the SS with one-hot-encoding but no significant improvement was observed when combining with the feature of OH+RNA-MSM_Emb. This result may be because the RNA-MSM model implicitly contains information on the secondary structure. Therefore, we did not include predicted SS as a feature for our final model.

After this work was completed, we noticed a new method called M^2^pred published on RNA solvent accessibility prediction, which utilized multi-scale context features via multi-shot neural network^55^. We downloaded this evolution-profile-based method from https://github.com/XueQiangFan/M2pred/ and compared it with RNA-MSM. The result shows that M^2^pred performs worse than our method (OH+RNA-MSM_Emb) not only on the newly released structures in the PDB (TS2) but also on the TS*, a subset of TS which removed similar structures to the sequences (TM-Score>0.45, according to RNA-align) in the training and validating set from their benchmark dataset. For this reduced set (TS*), the average correlation coefficient is 0.457 by M^2^pred and 0.504 by OH+RNA-MSM_Emb. The MAE is 34.66 by M^2^pred and 33.51 by OH+RNA-MSM_Emb. OH+RNA-MSM_Emb also shows a better distribution on TS2 and TS* in Supplement Figure S5. Thus, OH+RNA-MSM_Emb retains the best performance, when compared to the latest method.

One common problem of RNA language models including RNA-MSM is a limited sequence length due to the limited memory capacity of GPUs. The common reason for the high memory consumption of these RNA language models is because the transformer requires a space of magnitude in square of the input length. Even though we employed relatively new GPUs V100, the sequence length of the input can only be set to 1024. This leads to the loss of relationships between long distance nucleotides. The next version of the model should allow a better handing of sequence of arbitrary length using a method such as asymmetric cross-attention^56^.

One pressing problem is whether deep learning models can be generalized to the families that were not in the training or validation set^57^. We have made our best effort to separate training from validation and test by excluding structurally similar RNAs based on RNA-align. We further showed that the accuracy of models based on RNA-MSM features for several Rfam families unseen by the secondary structure predictor and the ASA predictor during training are as high as other RNAs. This suggests the model proposed here is able to go beyond the families contained in the training.

## Materials and Methods

### MSA generation

We downloaded 4069 RNA families (version 14.7) from https://rfam.xfam.org on 09/04/2022. The fully automatic RNAcmap3 for homolog search and sequence alignment^34^ was employed for these 4069 RNA families by using their covariance models (CM) for each family. Although the language model is unsupervised learning, we excluded the Rfam families which contains RNA sequences with experimentally determined structures to minimize potential over-fitting for structural inference. This leads to a total of 3932 Rfam families. The median value for the number of MSA sequences for these families by RNAcmap3 are 2184. All sequences were pre-processed by replacing ‘T’s with ‘U’s in RNA sequences and substituting ‘R’, ‘Y’, ‘K’, ‘M’, ‘S’, ‘W’, ‘B’, ‘D’, ‘H’, ‘V’, ‘N’ with ‘X’, similar to previous work^20^. The final vocabulary of our model contains of six letters: ‘A’, ‘G’, ‘C’, ‘U’, ‘X’, ‘-’. This dataset is called TR0 for unsupervised learning.

### Training and test sets for downstream models

To perform downstream tasks, we prepared two data sets based on experimentally determined RNA structures, which have the gold-standard information for base pairs and solvent accessibility. All RNA-containing structures were initially downloaded from the PDB^58^ website on August 9, 2021. They were randomly split into training (TR1, 80%), validation (VL1, 10%), and independent test (TS1, 10%), respectively. Sequences with similarity of more than 80% in VL1, TS1 were first removed using CD-HIT-EST^59^ and then, any sequences in TR1 with similar structures to the sequences in VL1 and TS1 (TM-Score>0.45, according to RNA-align^60^) were also removed. Similarly, any sequences in VL1 having similar structures with sequences in TS1 were also removed. Finally, we obtained 405 RNAs for TR1, 40 RNAs for VL1 and 39 RNAs for TS1. We updated the test set by downloading RNA structures deposited between 9, August 2021 and 14, July 2022 from the PDB website. We removed sequences by using CD-HIT-EST (80%) and RNA-align (TM-score of 0.45) against TR1, VL1, and TS1. This led to 31 RNA tertiary structures for TS2. The base pairs of all datasets (TR1, VL1, TS1, and TS2) were derived from their respective tertiary structures by using DSSR software^61^. We combined TS1 and TS2 to make the final independent test set with 70 RNAs (TS).

### Network architecture

As shown in Figure 1, RNA-MSM mainly consists of two modules: embedding and MSA transformer as in the protein MSA transformer work^24^. The embedding module consists of one initial embedding layer and two learnable position-embedding layers (Figure 1). The position-embedding layers encode rows (number of entries in the MSA) and columns (sequence length) of the MSA, separately. A 1D sequence-position embedding is applied to the row of the MSA, allowing the model to recognize the sequential order of nucleotides. In addition, each column of the MSA is embedded with a separate positional embedding, allowing the model to perceive the entire MSA as a series of ordered sequences.

We employed a similar configuration of MSA transformer module (Figure 1) as the protein MSA transformer work^24^. Briefly, this module is made of a stack of MSA transformer blocks. Each MSA transformer block has a residue and sequence attention layer with 12 attention heads with embedding size of 768, followed by a feedforward layer. LayerNorm is applied either before or after the attention layer. The residue attention layer captures interactions between nucleotides and integrates all the sequence attention maps within the MSA, resulting in single attention map being shared by all sequences to reduce memory usage. Moreover, sharing one attention map among all input sequences might learn the inherent structural information^24^. Through the feedforward layer, the input is passed over fully connected layers and activated using GELU activation functions. Specially, we modified a few specific parameters as follows. The number of blocks was changed from 12 to 10 due to the GPU memory limitation. We also changed the training precision from half-precision to 32-bit precision, which demonstrated an improved performance.

We define an input MSA as a matrix, *N* ×*L* where *N* represents the number of entries in the MSA and *L* represents the sequence length. After the embedding module, it was embedded into a tensor *N* ×*L*×768 and fed into the MSA transformer module. The final output contains two features: an embedding of *N* ×*L*×768 that represents the last MSA transformer block’s output and an attention map *L* ×*L*×120 derived from all residue attention layers where 120 was from multiplication of the number of attention heads (12) by the number of MSA transformer blocks (10).

### RNA-MSM Model training and inference

RNA-MSM was trained with a probability of 0.1 dropout after each layer. A total of 300 epochs were trained using eight 32G GTX V100 GPUs with Adam optimizer set to 0.0003, warmup step set to 16000, weight decay set to 0.0003, batch size set to 1, and learning rate set to 0.0003. The training stage stopped when the F1 value on validation set VL1 does not increase in 10 consecutive epochs.

For each input MSA, 1024 RNAs are randomly selected in addition to the query sequence if the number of sequences of input MSA is larger than 1024. It should be noted that the representative sequence is picked every time.

The maximum number of tokens is set to 16384 because the memory limitation of 32G of each V100 GPU. We employed the BERT masked language modeling objective function for training. Briefly, the token of the input MSA is masked by 20% at random. A total of 80% of these tokens are replaced with <MASK> token, 10% with random tokens, and the remaining 10% remain unchanged. The model is trained to predict the original tokens of the masked ones based on those tokens that are not masked in the sequence. During the inference, we employ the hhfilter^62^ to sample the RNA sequences in the maximum diversity manner, as the number of RNA sequences in some MSAs is huge. We experimented with the number of sampled sequences from all MSAs and found that sampling 512 sequences is a good balance of performance and computational cost.

### RNA secondary structure prediction

To establish a downstream task for RNA-MSM, we employed a simple ResNet^63^ similar to that utilized in (Singh, et al., 2019) ^38^. As shown in Supplementary Figure S1, the architecture of the model consists of 16 residual blocks followed by a fully connected (FC) block. Each residual block consists of two convolutional layers with a kernel size of 3×3 and 5×5 and with a filter size of 48. We tested one-hot-encoding, the embeddings as well as the attention map obtained from RNA-MSM as the input for the ResNet16 model. These input embeddings were converted into a three dimensional tensor by the outer concatenation function as described in RaptorX-Contact^64^. The model was implemented in Pytorch framework and use Nvidia graphics processing unit (GPU) to speed training. The model trained by using the Adam optimization function with a learning rate of 0.001 and the cross entropy was used as loss function. We trained the model on TR1 and the trained model was chosen based on the performance on VL1.

### RNA solvent accessibility prediction

Solvent accessible surface area (ASA) reflects the extent that a nucleotide in an RNA chain is exposed to solvents or other functional biomolecules. Unlike secondary structure dominated by local interactions, solvent accessibility is a measure of three-dimensional structure in one dimension. The ASA labels of TR1, VL1, and TS are calculated from their 3D structures by POPS package^65^ with a probe radius of 1.4 Å. All ASAs were normalized to relative accessible surface areas (RSAs). To be specific, we divided the ASA values by the maximum ASA of each corresponding nucleotide (i.e. A, G=400 Å^2^, U, C=350 Å^2^) as mentioned in (Yang et al., 2017)^43^.

We employed a simple network based on ResNet architecture to construct the RNA solvent accessibility prediction model. As shown in Supplementary Figure S6, this network mainly contains two blocks. The first block contains two convolutional layers^66^ to capture high-level feature maps and one standard “Squeeze- and -Excitation” (SE) module^67^, which defines models interdependencies between channels. The second block contains one self-attention layer^68^ and a simple multilayer perceptron layer (MLP). The self-attention layer is a core module in transformer architectures^68^, which jointly attend to different representation subspaces, and the multilayer perceptron layer fuse all the features and generates the outputs.

The outputs yielded by the model are values ranging from 0 to 1 as predicted relative solvent accessibility (RSA). The predicted ASA values were obtained by converting the RSA values to actual ASAs with the above-mentioned normalization factors. The mean absolute error between the predicted RSA and the actual RSA was used as a loss function. Model was trained on TR1 and validated on VL1. During the training, the learning rate scheduler of ‘Cosine Annealing Warm Restarts’^69^ was used to adjust the learning rate. The training iteration stops when the loss on the VL1 no longer decreases.

We determined the hyperparameters (number of repetitions for each type of residual block =1, width of the network = 64, learning rate = 0.005, batch size = 16, head of self-attention layers = 8) of the network by optimizing the performance on the VL1 dataset using one-hot encoding. After designing the network, we performed a number of experiments to examine whether the language embedding can improve the representations of sequence in ASA prediction. Different inputs for ASA prediction model, including the same sequence profile as RNAsnap2, one-hot encoding, embeddings and attentions from the language models, were tested.

## Code availability

All source codes, models, and datasets of RNA-MSM along with the RNA secondary structure predictor and ASA predictor are publicly available through https://github.com/yikunpku/RNA-MSM. We also built a freely accessible web server for using the RNA-MSM models, which is available at https://aigene.cloudbrain2.pcl.ac.cn/#/rna-msm.

## Author Contributions

These authors contributed equally to this work (Yikun Zhang, Mei Lang, Jiuhong Jiang, Zhiqiang Gao). Yaoqi Zhou, Jie Chen and Jian Zhan conceived of and supervised the study. Yikun Zhang, Mei Lang, Jiuhong Jiang and Zhiqiang Gao implemented the algorithms, performed the data analysis. Fan Xu, Thomas Litfin, Ke Chen, Jaswinder Singh, Xiansong Huang contributed to the protocol optimization and experimental design. Guoli Song, and Yonghong Tian provided scientific guidance and contributed to study supervision. Yaoqi Zhou, Yikun Zhang, Mei Lang, Jiuhong Jiang, and Zhiqiang Gao wrote the manuscript and all others have read and contributed the editing of and made the final approval of the manuscript.

## Notes

The authors declare no competing financial interest.

## Acknowledgements

We thank Shenzhen Science and Technology Program (Grant No. KQTD20170330155106581) and National Key Research and Development Program of China (NO.2021YFF1200400). The support of Shenzhen Bay supercomputing facility is also acknowledged.

